# Electroantennographic and behavioral responses of *Rhodnius prolixus* (Stal 1872) to xenobiotics reveal carvone and IR3535 as potential repellent candidates

**DOI:** 10.64898/2026.04.28.721475

**Authors:** Edwin Rodolfo Escobar Olarte, Gustavo Adolfo Rincón, María Fernanda Vidal, Ruth Mariela Castillo, Agustín Gongora, Sandra Carolina Montaño Contreras, María Carolina Velásquez-Martínez, Jonny Edward Duque

**Affiliations:** Centro de Investigaciones en Enfermedades Tropicales - CINTROP. Facultad de Salud. Escuela de Medicina, Departamento de Ciencias Básicas, Universidad Industrial de Santander, Guatiguará Technology and Research Park, Km 2 Vía El Refugio, Piedecuesta, Santander, Colombia; Grupo de Investigación en Neurociencias y Comportamiento UIS-UPB. Facultad de Salud, Escuela de Medicina, Departamento de Ciencias Básicas, Universidad Industrial de Santander, Bucaramanga, Santander, Colombia; Grupo de Investigación GYAS de la Facultad de Ciencias de la Salud. Escuela de Salud Pública, Universidad de los Llanos, Villavicencio, Meta, Colombia

**Keywords:** Olfactory response, repellent activity, insect behavior, screening system

## Abstract

Electroantennography (EAG) is a valuable approach for monitoring the sensory responses of insects to insecticidal and repellent molecules and an effective tool for early screening of compounds aimed at controlling and protecting against medically important insect vectors. However, its predictive potential for repellent efficacy in triatomine vectors remains poorly explored. The objective of this study was to evaluate the EAG responses to different xenobiotics as a preliminary selection strategy for compounds with potential repellent action against triatomines. For this purpose, the antennae of adult triatomines subjected to prolonged fasting (≥30 days) were exposed to repellent molecules. In parallel, repellency bioassays were conducted using a live bait (*Gallus gallus)* and a newly designed laboratory device to validate the electroantennographic results. EAG recordings showed a significant reduction in olfactory capacity of> 60% in response to the chemical compounds IR3535 and carvone, consistent with the protection times observed in the repellency tests (135.6 ± 43.29 min and 108 ± 26.33 min, respectively). In conclusion, the compounds with the highest repellent activity were clearly discriminated by the insects’ olfactory system, a finding corroborated by the decrease in electrical signals recorded in the EAG bioassays.

## Introduction

Chagas disease (CD) constitutes a significant public health problem in Latin America. It is caused by the flagellated protozoan *Trypanosoma cruzi*, transmitted mainly through contact with the feces of hematophagous insect vectors, commonly known as kissing bugs. These insects belong taxonomically to the family Reduviidae, subfamily Triatominae, comprising 158 species across 19 genera [1,2]. Because there is no single model of treatment, the most effective strategy to reduce its vector-borne incidence relies on the joint implementation of at least three preventive measures: housing improvement in affected areas, environmental education of communities in risk zones, and vector control through the use of focalized insecticide applications, taking into account the insect’s synanthropic behavior. This latter method is considered the most effective strategy to decrease the incidence of the disease, as reducing triatomine populations interrupts the transmission cycle. In line with this, commercial vector control products are mainly based on pyrethroids such as deltamethrin and cypermethrin. However, the intensive and prolonged use of these compounds over the last decades has led to the emergence of insect populations resistant to their active ingredients [3–5].

With approximately eight million people infected in Latin America and around one hundred million at risk of infection [1], it is paradoxical that repellent molecules have not been widely considered for their potential to protect against vector-borne transmission of this zoonotic disease. Active ingredients such as DEET, picaridin, and IR3535, as well as natural compounds like citronella (*Cymbopogon* spp), which are regarded as broad-spectrum repellents against hematophagous insects, have been questioned regarding their effectiveness against triatomines due to their limited protection and the lack of comparable evidence on host protection times [6–9]. A particularly noteworthy aspect is that DEET (N,N-diethyl-m-toluamide), a synthetic molecule considered the “gold standard” for protection against hematophagous insects such as *Aedes aegypti* (Linnaeus, 1762), is notably ineffective against *Rhodnius prolixus* (Stal, 1872). For this reason, it is essential to explore new repellent alternatives and investigate their modes and mechanisms of action further [9–11].

Given the urgent need to detect repellent molecules in triatomines and the lack of standardized methods to effectively discriminate compounds with this effect, the peripheral nervous system of insects has been proposed as a promising research alternative for the rational design of repellents. Specifically, studies have focused on proteins of the insect olfactory system, such as odorant-binding proteins (OBPs) and odorant receptors (ORs), which play a crucial role in insect behavior (Oliveira et al. 2018, 2017). For this reason, both OBPs and ORs constitute alternative molecular targets for identifying compounds with insecticidal or repellent activity, which can be applied to the control of Chagas disease vectors [12–14].

As mentioned previously, few studies have used the insect olfactory system as a behavioral assessment tool to select molecules with insecticidal or repellent activity for vector control and protection against bites. Some studies have conducted repellency bioassays through surface impregnation or toxicity assays based on a trial-and-error approach using different molecules under the hypothesis of insecticidal action (Cuadros et al., 2017; Hernández et al., 2010; Lutz et al., 2014; Sainz, et al, 2012; Penido & Schofield, 2000; Vassena & Picollo, 2003). However, to date, the responses of the triatomine olfactory system to insecticidal and repellent molecules have not been evaluated using electrophysiological techniques (May-Concha et al., 2018; Reisenman, 2014). The only existing report of this methodology concerns studies conducted on *Ae. aegypti*, focusing on natural compounds with repellent activity [15]. Therefore, the present study aimed to evaluate electroantennographic responses to discriminate xenobiotics with potential insecticidal and/or repellent activity in *R. prolixus*. The working hypothesis was that a significant reduction in electroantennography (EAG) responses to ammonia following exposure to a xenobiotic could predict its potential *in vivo* repellent effect.

## Materials and Methods

### Biological Material

Adult specimens of *R. prolixus* established for more than ten years in the insectary of CINTROP-UIS were used in this study. The insects were maintained in rearing rooms under controlled conditions of 26 ± 3 °C, 75 ± 4% relative humidity, and a 12:12 h light-dark photoperiod. Adult *R. prolixus* were fed every fifteen days with chicken blood (*Gallus gallus*) to obtain eggs and nymphs. For the electroantennography (EAG) experiments, the feeding interval was extended to achieve 30-day fasting periods, thereby stimulating antennal responsiveness.

### Molecules

To determine repellent activity, the molecular standards DEET (gold standard) and IR3535 were used, both of which are recognized as broad-spectrum insect repellents [6,8,15]. Ammonia (NH_3_) was employed as an attractant, while the test molecules consisted of the main constituents of essential oils previously reported to exhibit repellent activity against hematophagous insects [16,17]. Finally, acetone (99.98%, Merck ©) was used as a neutral control in both bioassays (Figure 1) (Geier, Bosch, and Boeckh 1999; Portilla Pulido et al. 2022).

**Figure 1.**
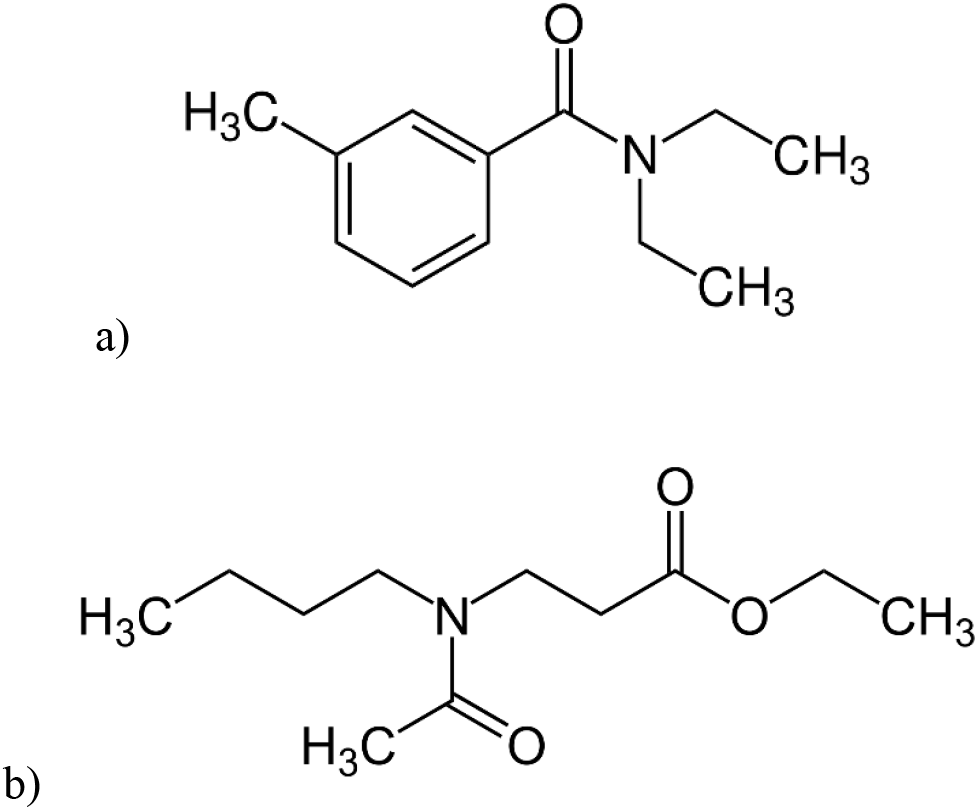
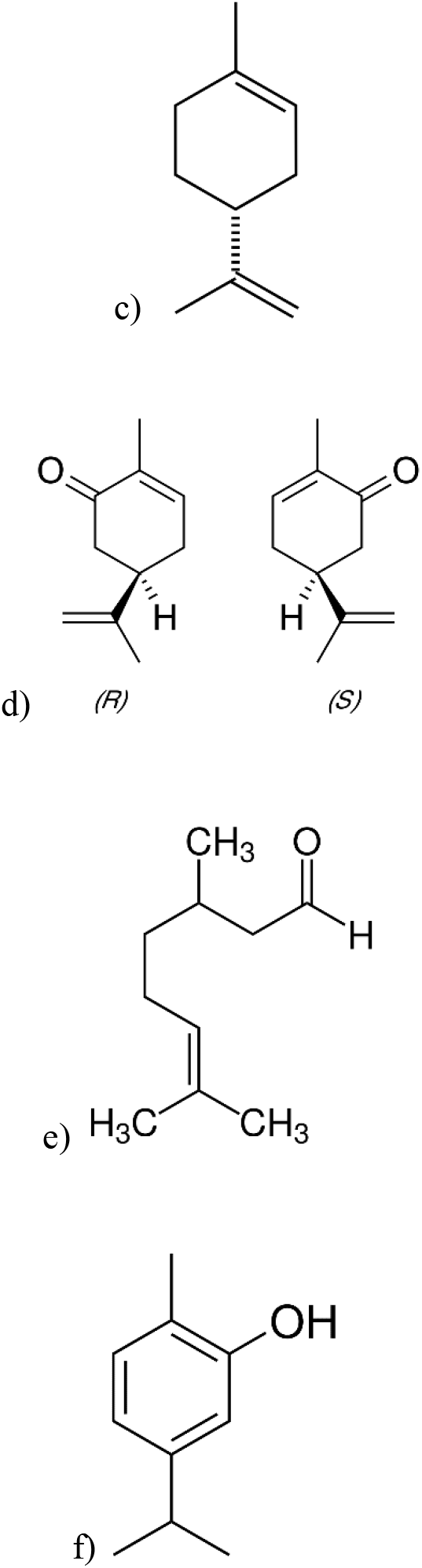
Repellent molecules used in the experiments: (a) DEET, (b) IR3535, (c) Limonene, (d) Carvone (R and S enantiomers), (e) Carvacrol, and (f) Citronellal.

### Insect Mounting for Electroantennographic Characterization

Adult *R. prolixus* individuals subjected to a 30-day programmed fasting period were used to evaluate the electroantennographic (EAG) response of the antennae upon exposure to the test molecules. Each insect was carefully positioned on a wooden platform and immobilized with modeling clay, leaving the antennae and eyes exposed (Figure 2). The mounted specimen was placed under a stereomicroscope (Leica EZ4) to facilitate the insertion of two tungsten microelectrodes (0.25 mm, Sigma-Aldrich): one serving as the recording electrode and the other as the reference. Before recordings, the electrodes were electrolytically sharpened by repeatedly immersing their tips in a 1 M sodium hydroxide (NaOH) solution (Sigma-Aldrich) at 3–5 V until an approximate tip diameter of 0.05–1 mm was reached, following the procedure described by Olsson and Hansson (2013). The reference electrode was inserted into the exposed eye, while the recording electrode was positioned at the base of the antennal pedicel. EAG signals were recorded two minutes after the insect was fixed to the mounting platform, amplified (1000×; Universal Single Probe, Type PRS-1, Syntech, Germany), digitized (IDAC-4, Syntech, Germany), visualized, and analyzed using Autospike software (Syntech, Germany) as previously described by [18].

**Figure 2.**
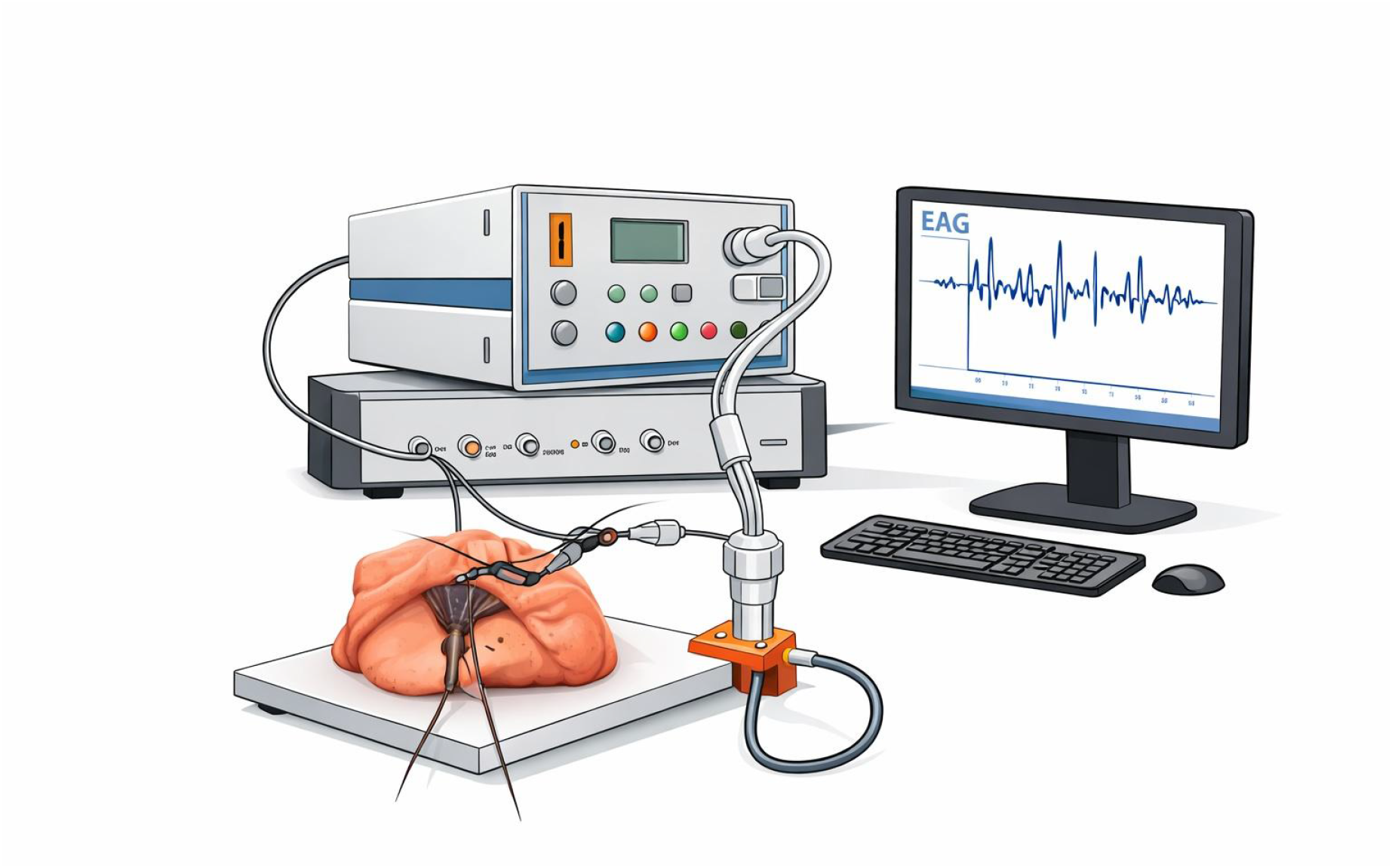
Electroantennography (EAG) experimental setup used to evaluate the olfactory responses of *Rhodnius prolixus* to volatile compounds. The insect was immobilized using modeling clay, leaving the head and antennae exposed for electrophysiological recordings. A recording electrode was inserted into the antenna, while a reference electrode was positioned in the eye region. Volatile stimuli were delivered through a controlled airflow system directed toward the antenna. Electrical signals generated by the antennal response were amplified with an EAG amplifier and recorded using a data acquisition system, enabling visualization and analysis of the electroantennographic signals.

### Olfactory stimulation and electroantennographic recording

Filter paper squares (0.5 × 0.5 cm, Whatman No. 1) were impregnated with each compound to be tested at different concentrations (30%, 60%, and 90%) and allowed to dry for one minute to ensure complete solvent evaporation [9,10]. Each impregnated square was then placed inside a glass test tube with a lateral outlet connected to the airflow controller. Pulses of air were delivered through the tube to the insect preparation via a controlled airflow pump (Stimulus Controller Unit, Type CS-55, Syntech, Germany). The airflow pump was programmed to emit a sequence of three air pulses at a flow rate of 25 mL/s, each lasting 10 seconds, with 2-second intervals between pulses containing the repellent compounds and 8 seconds of clean air serving as background airflow.

Before and after each repellent application, a 25% ammonia pulse was delivered to ensure the insect remained responsive to stimuli throughout the experiment and to monitor variations in the electroantennographic signal. One individual was used per dose, with initial and final ammonia stimulation and 8-second air pulses (background) between doses. Independent materials (test tubes and connecting tubing) were used to evaluate each compound, with one individual per replicate (N = 7 per experiment).

### Evaluation of Repellent Activity

To determine the repellency of the proposed xenobiotics, bioassays previously reported in the literature were used as references [9,10]. However, due to limited ergonomics and handling difficulties associated with those designs, a new device was developed by the Medical Entomology Laboratory at CINTROP, based on the two previously reported models. A series of comparative bioassays was conducted between the two device types to validate the accuracy and reliability of the newly designed apparatus for determining repellent activity.

### CINTROP Device Specifications

The device was constructed from 3 mm-thick transparent acrylic sheets. It has a rectangular shape, measuring 10 cm in length and 3.6 cm in width, providing a total contact area of 36 cm^2^ (Figure 3). The proximal end features a mesh and an additional acrylic component that allows the insertion of filter paper, enabling either direct or indirect contact with live bait. Located 3 cm from the mesh, an internal division engraved into the acrylic separates the Bait Zone from the Intermediate Zone. The distal end features a movable gate positioned 3.2 cm from the device’s farthest edge, which defines the Refuge Zone. The apparatus also includes a fixation system for use with live, non-anesthetized bait (e.g., a chicken).

**Figure 3.**
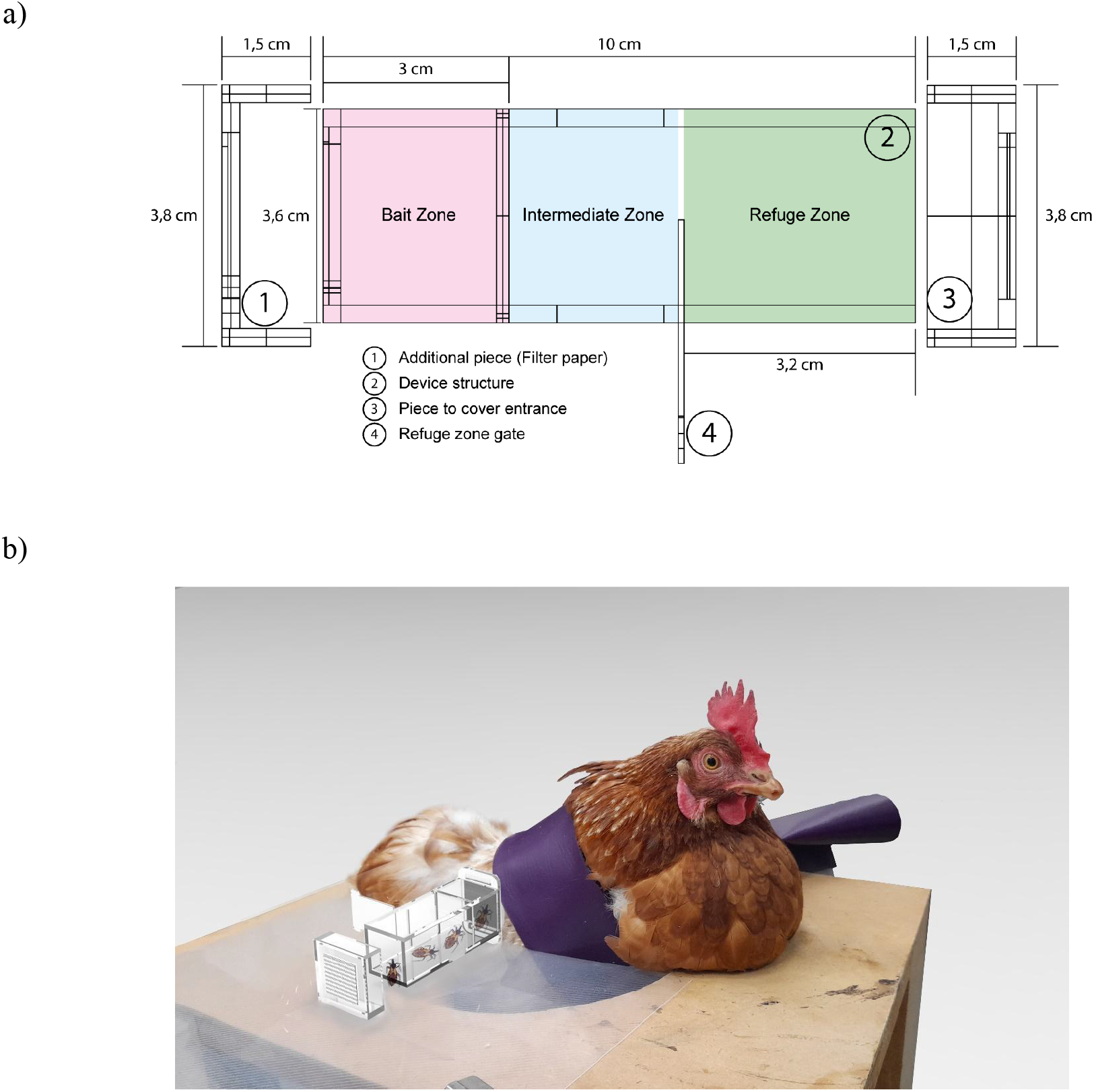
Design specifications of the proposed device for evaluating repellent activity in triatomines. (a) Schematic representation; (b) Device coupled to the bait.

### Determination of repellent activity

Both devices (Zermoglio and CINTROP) were evaluated simultaneously using adult chickens (*Gallus gallus*) as live bait and adult individuals of *R. prolixus*. Each bird was topically treated with 150 µL of the test solution applied to one side of the abdomen, beneath the wings. Initially, the commercial repellent IR3535 was tested at a concentration of 90%. The behavioral response of the insects was assessed using both the plastic device described by Zermoglio et al. [10] and the CINTROP prototype. For each molecule, three technical replicates were performed per day, with 15 insects per replicate, along with two control replicates (acetone). The entire experiment was repeated on different days using different live baits, constituting independent biological replicates.

Protection against biting (PC) and approach to the bait (AC) were recorded from 0 to 150 minutes, including 30-minute resting intervals to ensure animal welfare. Subsequently, the repellent effects of other xenobiotics with potential activity, including DEET, limonene, carvone, carvacrol, and citronellal, were evaluated at a concentration of 90%.

### Statistical Analysis

Electroantennographic (EAG) responses were calculated as the mean maximum amplitude (mV) recorded in response to each administered air pulse. All repellence and EAG data were tested for normality using the Kolmogorov–Smirnov test. Analysis of variance was then performed to compare maximum EAG responses among the different compounds (ANOVA or Kruskal–Wallis, depending on data normality), followed by multiple comparison tests (Tukey’s or Dunn’s, depending on the normality results). Only p-values < 0.05 were considered statistically significant.

## Results

### EAG records of xenobiotic compounds

To evaluate the olfactory response of *R. prolixus*, electroantennographic (EAG) signals were recorded after exposure to the tested xenobiotic compounds. Each insect exhibited a distinct EAG pattern, resulting in a wide range of amplitudes among individuals. Consequently, data normalization was performed using the initial and final ammonia amplitude responses from all tested compounds, taking the initial ammonia response as the 100% reference. A preliminary bioassay was conducted to assess the insects’ response to acetone, the solvent used for all tested compounds, and to confirm that it did not interfere with odorant detection. The results showed no statistically significant differences between the insects’ initial and final ammonia responses, indicating that acetone did not alter their olfactory perception. Therefore, it was confirmed that the solvent did not affect the activity of the compounds evaluated in the EAG assays (Figure S1).

Similarly, the same procedure was applied to compounds with potential repellent activity, which were dissolved in acetone at concentrations of 90%, 60%, and 30% (v/v). An initial ammonia recording was taken, followed by exposure to the repellent compound, and finally, a second ammonia recording was performed. The insects were able to perceive the odorants emitted by these compounds; however, the recorded responses were considerably lower than those elicited by ammonia, not exceeding 10% of its amplitude. Moreover, most compounds exhibited a response pattern similar to that of the negative control (acetone), as no statistically significant differences were observed between the initial and final ammonia recordings. Nevertheless, the compounds IR3535 and carvone showed a marked decrease greater than 60% in the final ammonia response, being the only two compounds that presented statistically significant (*Kruskal–Wallis* test, *H*(3, *N* = 21) = 17.44, *P* < 0.05; Dunn’s post hoc test) differences between their initial and final ammonia recordings (Figure 4)

**Figure 4.**
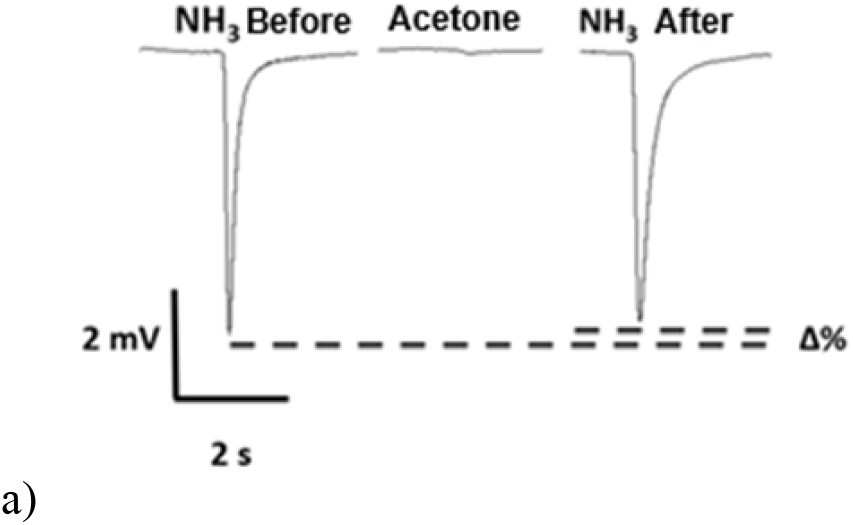
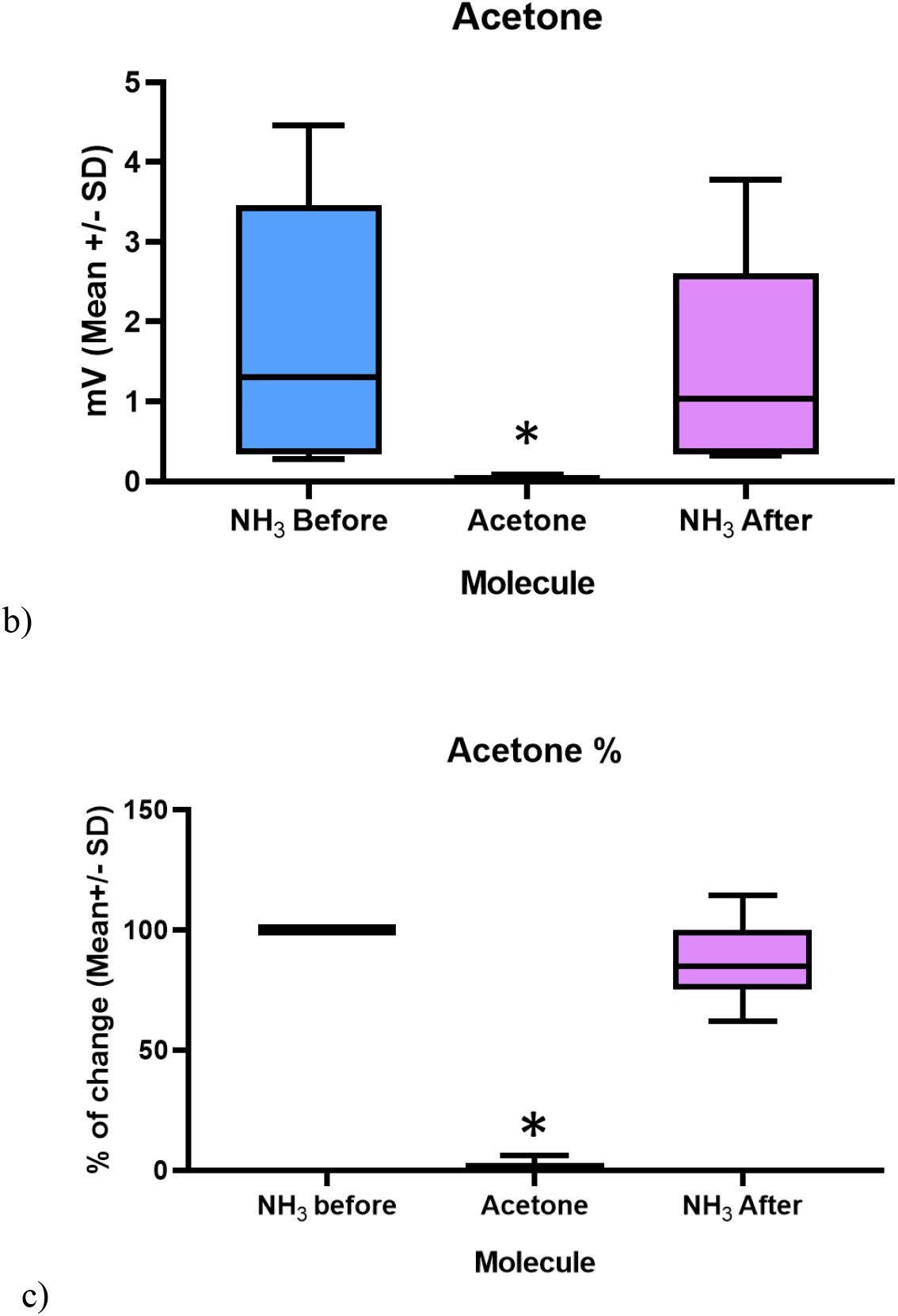
Electroantennographic (EAG) signals and percentage change in the ammonia response after exposure to acetone (negative control) with pre- and post-ammonia stimulation. (a) Typical EAG responses. (b) Electroantennographic signal recordings expressed in millivolts (mV, mean ± standard deviation). (c) Percentage change in the response to ammonia (mean ± standard deviation). No statistically significant differences were observed between the EAG signals to ammonia before and after acetone exposure (*Kruskal–Wallis* test, *H*(3, *N* = 21) = 17.44, *P* < 0.05; Dunn’s post hoc test).

When performing multiple comparisons of the EAG responses, it was observed that only the compounds IR3535 and carvone (one-way *ANOVA, F*(8, 45) = 37.43, *P* < 0.05; Tukey’s post hoc test, Figure 5) showed statistically significant differences when comparing the ammonia stimulus response before and after compound exposure. The remaining evaluated compounds did not exhibit statistically significant differences. They displayed a response pattern like that of the negative control (acetone), where the variation in the ammonia EAG signal showed no significant changes after compound application (*Kruskal–Wallis* test, *H*(3, *N* = 21) = 13.54, *P* > 0.05; Dunn’s post hoc test, Figure 4).

**Figure 5.**
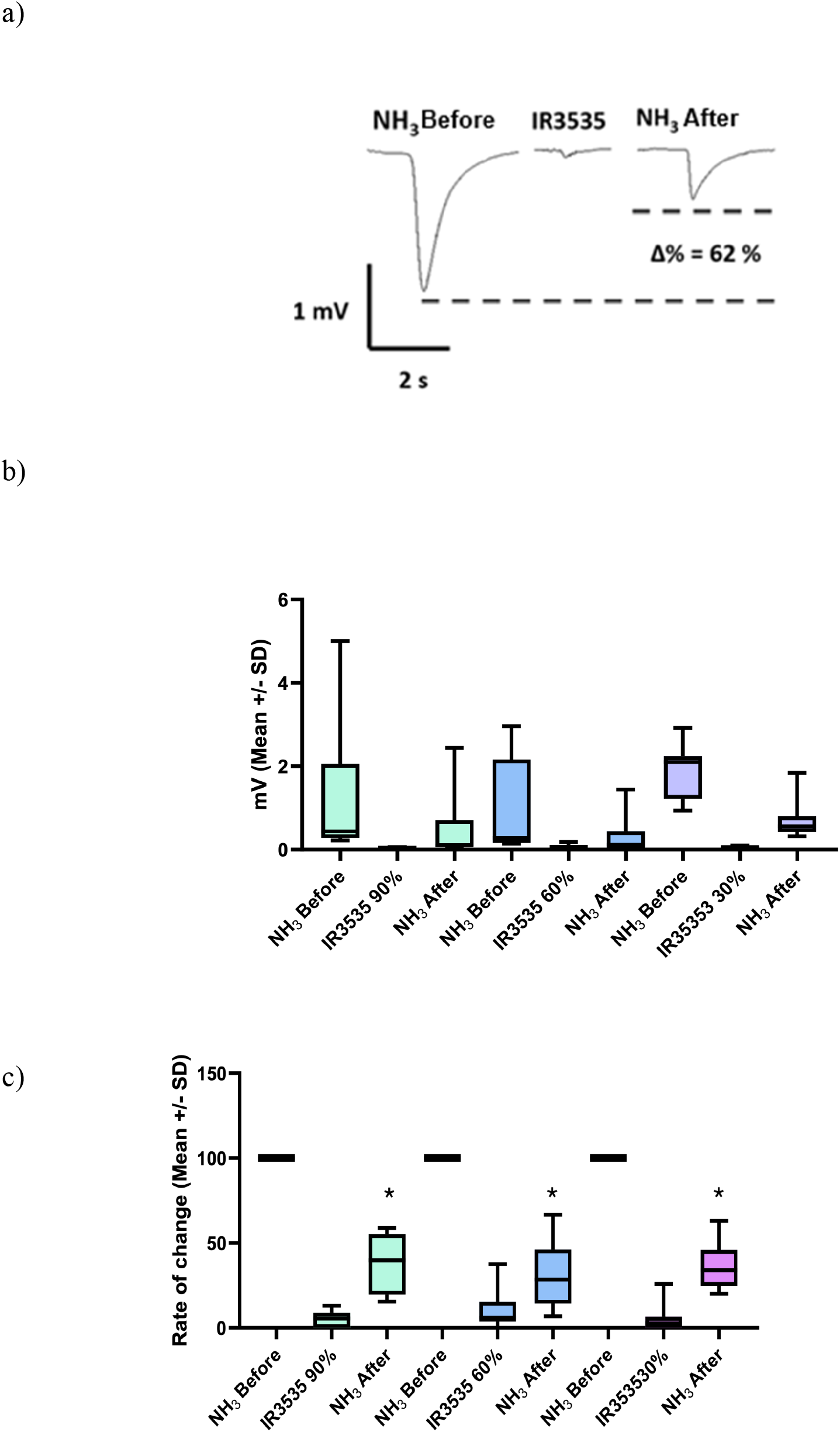
Electroantennographic signals and percentage change in the ammonia response after exposure to the IR3535 molecule (commercial repellent) using pre- and post-ammonia stimuli. a) Representative electroantennogram (EAG) responses. b) Electroantennographic signal recordings at three concentrations (90%, 60%, 30%) expressed in millivolts (mV), mean ± standard deviation. c) Percentage change in the response to ammonia at three concentrations (90%, 60%, 30%), mean ± standard deviation. (*) Indicates statistically significant differences between ammonia EAG signals before and after exposure to the molecule (Kruskal–Wallis test, H(3, N=21) = 17.44, *P* < 0.05; post hoc Dunn’s test).

**Figure 6.**
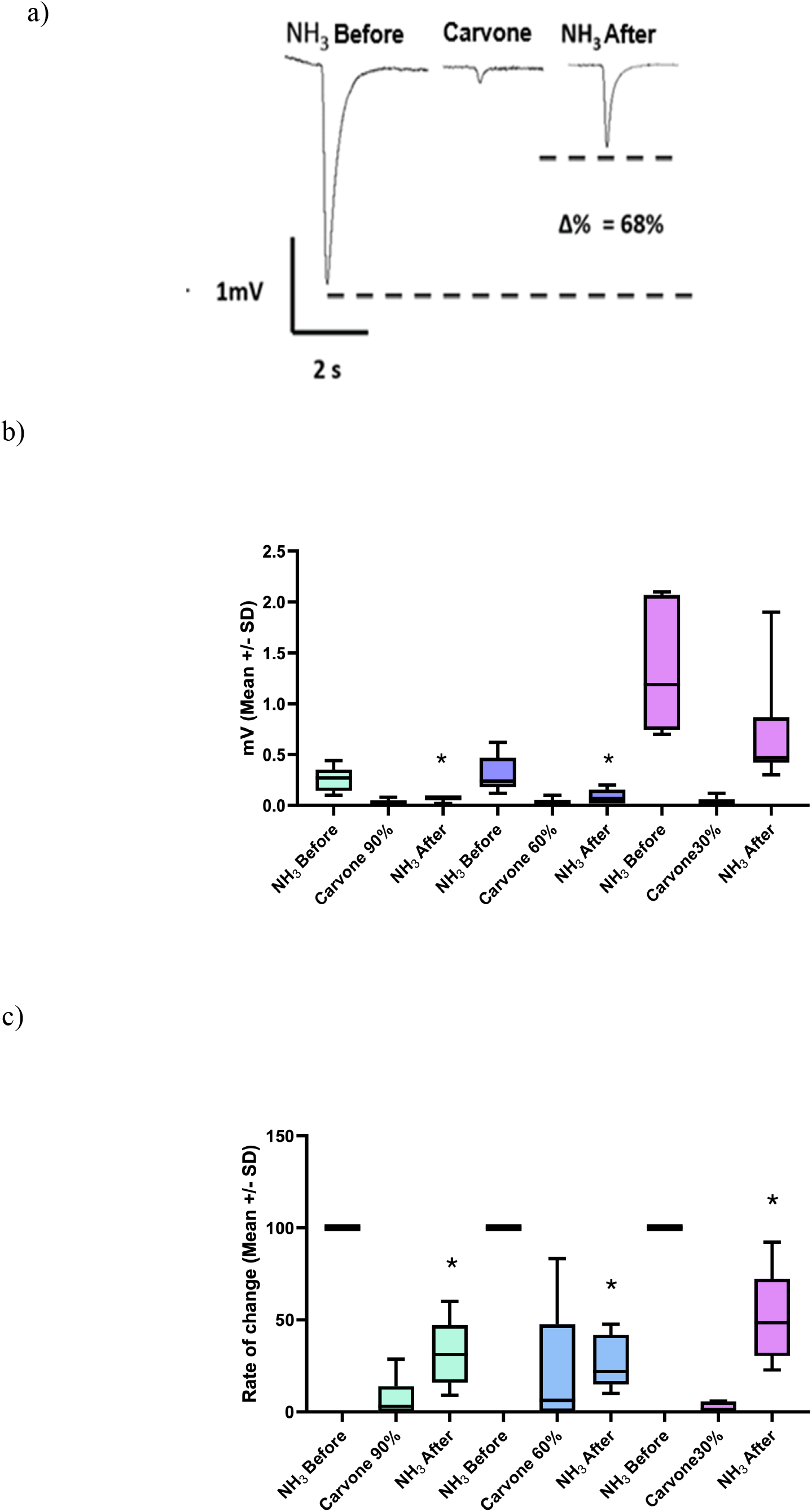
Electroantennographic signals and percentage change in the ammonia response after exposure to the carvone molecule, using pre- and post-ammonia stimuli. a) Representative electroantennogram (EAG) responses. b) Electroantennographic signal recordings at three concentrations (90%, 60%, 30%) expressed in millivolts (mV), mean ± standard deviation. c) Percentage change in the response to ammonia at three concentrations (90%, 60%, 30%; mean ± standard deviation). (*) Indicates statistically significant differences between ammonia EAG signals before and after exposure to the molecule (one-way ANOVA, F(8, 45) = 37.43, *P* < 0.05; post hoc Tukey test).

### Repellent Activity

#### Comparison between CINTROP and Zermoglio devices

In the comparison between the CINTROP and Zermoglio devices, protection against bites produced by the IR3535 compound at a 90% concentration was evaluated as a parameter of repellent activity. No statistically significant differences in protection time were observed between the two devices used against *R. prolixus* (Table 1). However, statistically significant differences were detected between the repellent compound IR3535 and its control (acetone) (Prueba U de Mann-Whitney, *P* < 0.05).

**Table 1.** Comparison of protection time between the Zermoglio and CINTROP devices. Mean protection time against *R. prolixus* bites (min ± SD) using IR3535 at 90% concentration.

| **Xenobiotics** | **Zermoglio**<br>**Time in minutes**<br>**(% ± SE)** | ***Cintrop***<br>**Time in minutes (%±SE)** |
| --- | --- | --- |
| **Negative control (Acetone)** | 12,18 ± 8,68<sup>a</sup> | 7,46 ± 6,97<sup>a</sup> |
| **IR3535 at 90%**<br>**concentration** | 118,43 ± 27,34<sup>b</sup> | 134,45 ± 26,93<sup>b</sup> |

*Identical letters indicate that there are no statistically significant differences between devices (Mann-Whitney U test, $U = 2.0$ , $p = 0.38$ ); different letters indicate significant differences between the repellent and its control within each device ( $P < 0.05$ ) in *R. prolixus*.*

Considering these results and the absence of statistically significant differences among the devices used in the standardization bioassays, the device designed by the CINTROP laboratory was selected for subsequent experiments. This device offers an adaptable design for the live bait used in the assays (chicken), preventing disturbances from bait movement and thereby ensuring proper completion and reproducibility of the bioassays.

#### Evaluation of xenobiotics with repellent activity

Two commercial repellents (DEET and IR3535) and three major components of essential oils (limonene, carvone, and carvacrol) were evaluated at 90% concentration. Acetone was used as a negative control. The compound showing the highest repellent activity was IR3535, with a protection time against insect bites of 135.6 ± 43.29 minutes (mean ± standard deviation), followed by carvone, with a mean protection time of 108 ± 26.33 minutes. Both compounds showed statistically significant differences compared with the other molecules evaluated, with protection lasting at most 5 minutes. Statistical analysis was performed using the Kruskal–Wallis test (H(7, N = 68) = 37.12, P < 0.05) followed by Dunn’s post hoc test (Figure 7).

**Figure 7.**
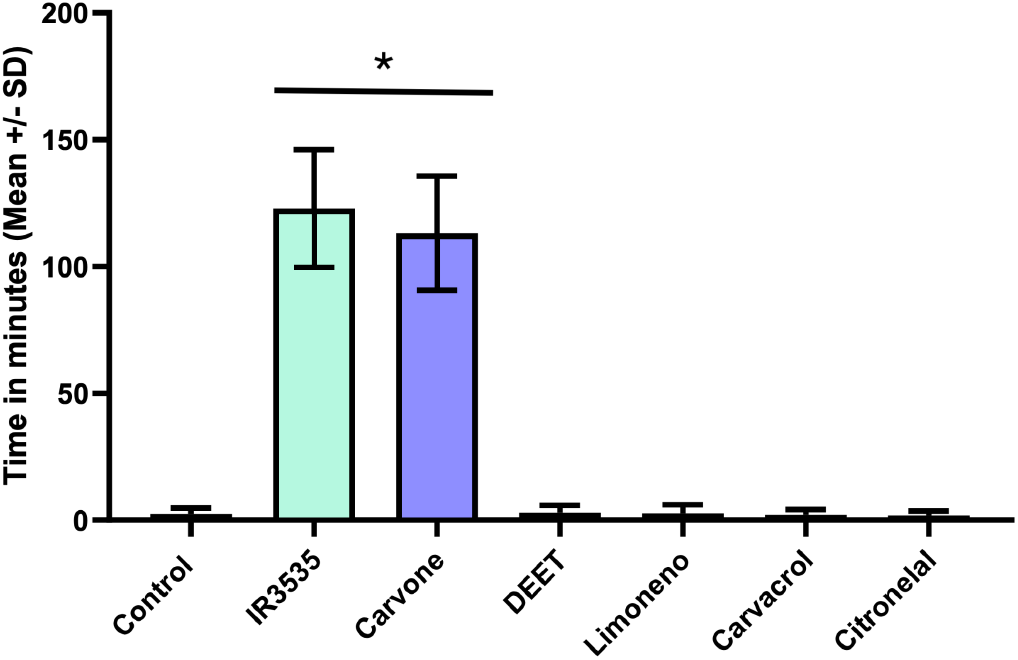
Protection times (mean ± SD) against insect bites for compounds with potential repellent activity: IR3535, carvone, DEET, limonene, carvacrol, citronellal, and acetone (control), all tested at 90% concentration. Differences among treatments were evaluated using the Kruskal–Wallis test (H(7, N = 68) = 37.12, P < 0.05), followed by Dunn’s post hoc test. (*) Indicates statistically significant differences among the evaluated compounds.

#### Approach to the host

In addition to protection time, the host-approach time was recorded as the first movement of the insect toward its feeding source, representing the initial intention to bite the host, regardless of whether the feeding attempt was successful. In the presence of molecules with potential repellent activity, insects approached the host but, under the influence of the compound, retreated to the refuge zone, where they performed multiple attempts to approach. For behavioral analysis, the first biting attempt was used as the reference point and expressed in minutes (Table 2). The recorded times were comparable to the protection times obtained in the repellent assays. A similar pattern was observed in compounds without repellent activity; however, in those cases, puncture wounds were observed on the chicken skin, whereas no lesions were detected when repellent compounds were tested. No statistically significant differences were observed among the bioassays performed (ANOVA, F(6,61) = 1.066, P = 0.39).

**Table 2.**
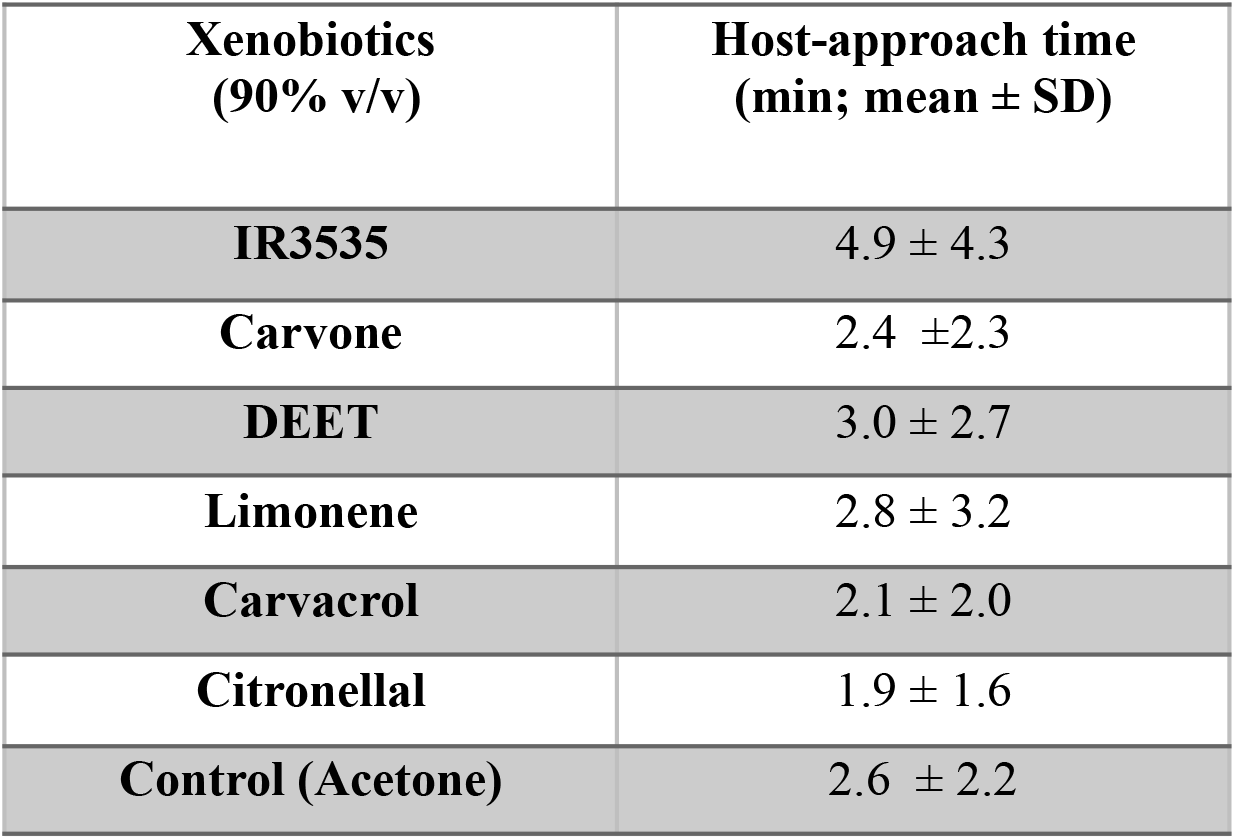
Host-approach time for the evaluated compounds: IR3535, carvone, DEET, limonene, carvacrol, citronellal, and control (mean ± standard deviation). No statistically significant differences were observed among the bioassays performed (ANOVA, F(6,61) = 1.066; P = 0.39).

| **Xenobiotics**<br>**(90% v/v)** | **Host-approach time**<br>**(min; mean ± SD)** |
| --- | --- |
| **IR3535** | 4.9 ± 4.3 |
| **Carvone** | 2.4 ± 2.3 |
| **DEET** | 3.0 ± 2.7 |
| **Limonene** | 2.8 ± 3.2 |
| **Carvacrol** | 2.1 ± 2.0 |
| **Citronellal** | 1.9 ± 1.6 |
| **Control (Acetone)** | 2.6 ± 2.2 |

## Discusion

### Electroantennographic recordings

The present study focuses on the characterization of olfactory chemoreception in triatomines to establish a quantitative correlation between biological activity (repellency/attraction bioassays) and peripheral electrophysiological responses (electroantennographic recordings, EAG) of these vectors when exposed to a series of volatile compounds. This approach aims to develop an early screening system for discriminating repellent molecules based on the insect’s physiological responses.

Initially, the olfactory system of insects was stimulated with ammonia (NH_3_), a compound known to play a key role in host-seeking behavior in hematophagous insects and is commonly used as a standard stimulus in electroantennographic (EAG) assays due to its consistent, concentration-dependent responses in *R. prolixus* [19]. As expected, stimulation with ammonia elicited clear electrophysiological responses in the olfactory neurons, with signal amplitudes ranging from 0.26 to 4 mV. These responses reflect activation of ion channels involved in odor reception, with higher voltage amplitudes indicating greater ionic flux across neuronal membranes, resulting in membrane depolarization [20].

However, the responses elicited by the evaluated compounds were considerably lower than those produced by ammonia. This difference may be explained by these compounds activating fewer olfactory receptors or ion channels, as reflected in the reduced amplitude of the recorded signals [20]. In this context, ammonia was subsequently used again as a reference stimulus to evaluate how the tested compounds, acting as repellents or insecticides, modulate the olfactory system’s response to this attractant.

### Responses to repellent compounds

The present results indicate that several volatile compounds with repellent activity can modulate the olfactory responses of *R. prolixus*, as evidenced by both behavioral assays and electroantennographic (EAG) recordings. In particular, compounds such as IR3535 and carvone produced measurable effects on the insects’ antennal responses, as confirmed by comparing initial and final ammonia recordings, in which a reduction of approximately 60% in signal amplitude was observed at the highest dose. This decrease suggests that these molecules may interfere with the insect’s ability to detect host-related cues, thereby affecting host-seeking behavior. From a physiological perspective, reductions in EAG amplitude are generally associated with changes in the activity of odorant receptor neurons (ORNs), which play a central role in olfactory perception in hematophagous insects. Similar effects have been described for well-known repellents such as DEET, which can attenuate antennal responses to attractive odors by inhibiting or modulating ORN activity [21]. Other compounds, including picaridin and IR3535, have also been reported to affect olfactory signal transduction in blood-feeding insects through comparable mechanisms [22]. Together, these findings support the idea that repellent compounds may disrupt host detection by altering peripheral olfactory signaling, highlighting the usefulness of EAG recordings as a screening tool for identifying molecules with potential repellent activity against vectors of Chagas disease.

### Molecular mechanism of repellent compounds

The molecular mode of action of repellent compounds has been proposed to involve the selective inhibition of heteromeric odorant receptor (OR) complexes, which are insect-specific chemosensory proteins responsible for detecting airborne odorants. However, this model remains controversial, as some studies have shown that repellent compounds can both activate and inhibit odorant receptor neurons in adult insects, ultimately triggering avoidance behavior. Taken together, these findings suggest that repellents may interact with multiple biological targets distributed across different chemosensory pathways [21,23]. In this context, a compound may be considered to exhibit repellent activity when EAG assays show a statistically significant reduction in the electrical response to ammonia after exposure to the test molecule. In the present study, this pattern was observed for IR3535 and carvone, which produced approximately a 60% decrease in signal amplitude and also showed protective effects in behavioral assays, with protection times exceeding 100 minutes in live-bait repellent tests.

In contrast, compounds that provided protection times shorter than 5 minutes did not produce statistically significant changes in the electroantennographic responses to ammonia. The similarity between the initial and final ammonia recordings indicates that ammonia does not significantly affect the insects’ olfactory system and therefore does not exhibit repellent activity. Evidence supporting the importance of olfactory pathways in host detection has been reported in triatomines in which odorant receptors and the co-receptor RproOrco were silenced using RNA interference (RNAi). In those studies, a 73% reduction in this gene’s expression resulted in a marked impairment in the insects’ ability to locate a vertebrate host [24]. These findings highlight the central role of odorant receptor complexes in host-seeking behavior and support the interpretation that compounds such as IR3535 and carvone may exert their repellent effects by interfering with these chemosensory proteins. This interpretation is consistent with the results of the present study, in which both compounds significantly reduced ammonia-induced electrical signals in EAG assays and showed protective effects in behavioral bioassays. Taken together, these findings suggest that electroantennographic assays may serve as an early screening approach to predict the repellent potential of unknown compounds. In this context, electroantennography emerges as a complementary and predictive tool for screening repellent molecules against triatomines, thereby strengthening non-insecticidal strategies for vector control of Chagas disease.

### Repellent activity

As discussed in the background section on devices used to measure repellent activity, there is currently no fully standardized methodology for estimating this variable in triatomines. Among the available approaches, arena tests combined with motion tracking systems (e.g., Videomex) are commonly used [8]. However, this methodology has important limitations, particularly the absence of a live host stimulus, which makes it difficult to evaluate the insect’s actual biting intention. Another common feature of these assays is the use of DEET as a reference compound to demonstrate repellent activity. Nevertheless, both arena assays and live-bait devices indicate that high concentrations of DEET are required to observe a measurable repellent effect. In live-bait bioassays, concentrations as high as 90% have been reported [10], and some studies have even suggested limited efficacy of DEET against *R. prolixus* [25].

For this reason, IR3535 was selected as the reference compound to evaluate repellent activity in the present study. A 90% concentration was used, and a new device was designed to assess repellent activity, based on the device originally proposed by Zermoglio, while maintaining its main structural characteristics. When comparing the Zermoglio and CINTROP devices, no statistically significant differences were observed in the protection time obtained with IR3535. These results indicate that repellent activity can be reliably measured using either device. However, the device developed by the Medical Entomology Laboratory (CINTROP) offers several practical advantages, including improved ergonomics, easier handling, adaptability to different animal baits via adjustable straps, and the ability to conduct multiple behavioral observations, as described in previous studies [9,10].

### Repellent activity of tested compounds

Among all the compounds evaluated, only the xenobiotic carvone showed a clear protective effect against insect bites, providing protection times exceeding 100 minutes. This response was statistically similar to that observed for the reference compound IR3535. In contrast, both compounds showed significant differences compared with the remaining tested molecules, which did not exceed 5 minutes of protection. These results indicate that only IR3535 and carvone exhibited measurable repellent activity under the experimental conditions used in this study [26].

## Conclusion

Overall, the results obtained in this study demonstrate that carvone and IR3535 exhibit measurable repellent activity against *R. prolixus*, as evidenced by both behavioral assays and electroantennographic responses. The integration of electrophysiological recordings with live-bait behavioral bioassays enabled the identification of compounds that interfere with the olfactory perception of host-related cues in this vector. In addition, the evaluation and validation of the CINTROP device provide a practical methodological alternative for measuring protection time under conditions that better simulate natural host–vector interactions. Taken together, these findings support the use of combined behavioral and electrophysiological approaches as effective tools for early screening of repellent compounds and highlight the potential of such strategies for developing non-insecticidal interventions to improve vector control of Chagas disease.

## Acknowledgments

This work was supported by the Ministerio de Ciencia, Tecnología e Innovación (MinCiencias), through the Fondo Nacional de Financiamiento para la Ciencia, la Tecnología e Innovación – Fondo Francisco José de Caldas, under grant numbers 833-2018 and 891-2020.

