## Supplemental material for "Electroantennographic and behavioral responses of *Rhodnius prolixus* (Stal 1872) to xenobiotics reveal carvone and IR3535 as potential repellent candidates"

a)

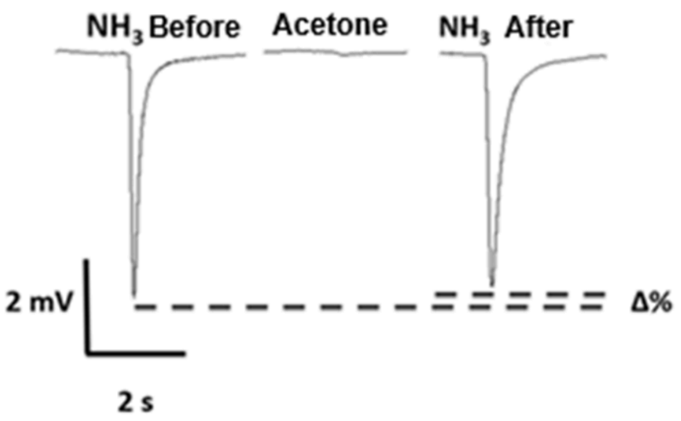
NH3 Before
Acetone
NH3 After
2 mV
2 s
Δ%

b)

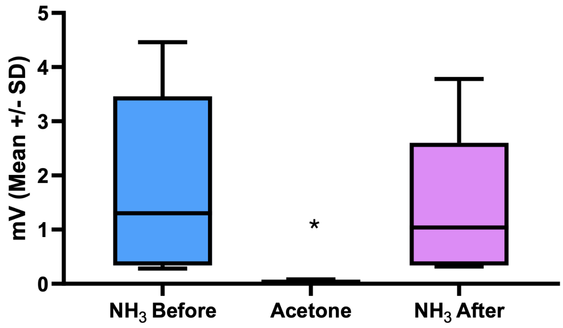
mV (Mean +/- SD)
5
4
3
2
1
0
*
NH3 Before
Acetone
NH3 After

c)

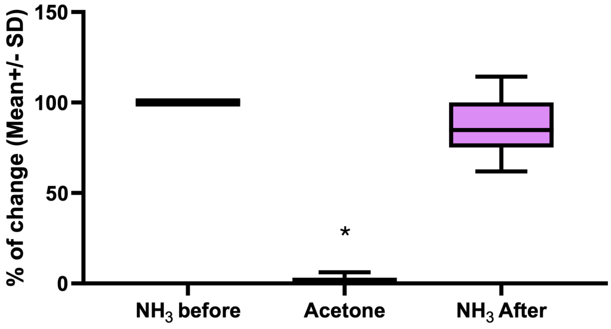
% of change (Mean+/- SD)
150
100
50
0
*
NH3 before
Acetone
NH3 After

Fig Supplementary 1. *Electroantennographic (EAG) signals and percentage change in the ammonia response after exposure to acetone (negative control) with pre- and post-ammonia stimulation.* (a) Typical EAG responses. (b) Electroantennographic signal recordings expressed in millivolts (mV, mean ± standard deviation). (c) Percentage change in the response to ammonia (mean ± standard deviation). No statistically significant differences were observed between the EAG signals to ammonia before and after acetone exposure (*Kruskal–Wallis* test, $H(3, N = 21) = 17.44$, $P < 0.05$; Dunn's post hoc test).
